# A Data-Dependent Acquisition Ladder for Ultrasensitive (Neuro)Proteomics

**DOI:** 10.1101/2021.08.03.454943

**Authors:** Sam B. Choi, Pablo Muñoz-LLancao, M. Chiara Manzini, Peter Nemes

## Abstract

Measurement of broad types of proteins from a small number of cells to single cells would help to better understand the nervous system but requires significant leaps in high-resolution mass spectrometry (HRMS) sensitivity. Microanalytical capillary electrophoresis electrospray ionization (μCE-ESI) offers a path to ultrasensitive proteomics by integrating scalability with sensitivity. We report here a data acquisition strategy that expands the detectable and quantifiable proteome in trace amounts of digests using μCE-ESI-HRMS. Data-dependent acquisition (DDA) was programmed to progressively exclude high-intensity peptide signals during repeated measurements. These nested experiments formed rungs of our “DDA ladder.” The method was tested for replicates analyzing ~500 pg of protein digest from cultured hippocampal (primary) neurons (mouse), which estimates to the total amount of protein from a single neuron. Analysis of net amounts approximating to ~10 neurons identified 428 nonredundant proteins (415 quantified), an ~35% increase over traditional DDA. The identified proteins were enriched in neuronal marker genes and molecular pathways of neurobiological importance. The DDA ladder deepens the detectable proteome from trace amounts of proteins, expanding the analytical toolbox of neuroscience.

## INTRODUCTION

Ultrasensitive high-resolution mass spectrometry (HRMS) presents a window to the molecular composition of cells, raising a potential to better understand neurobiological processes at the level of neurons and neural circuits. Using high-performance nano-flow liquid chromatography (nanoLC) HRMS, it is now possible to quantify ~13,000 different proteins from neuron cultures.^1-2^ To deepen the detectable coverage of the proteome, shot gun proteomics currently processes sizable populations, usually millions, of neurons to analyze ~100 ng–1 μg of protein. Averaging over an entire cell population, however, potentially loses information critical to particular neurons or neural circuits; it also elevates chemical noise during measurements, further limiting insights into the biochemistry of the neuron.^3^ A single mammalian neuron yields only ~500 pg of protein extract,^4^ which is ~1000–100,000-fold less than usually detected in nanoLC HRMS. Identification and quantification of the single-neuron proteome hinges on the development of HRMS approaches capable of trace-level sensitivity.

Ultrasensitive HRMS bridges over this technological gap for the proteome (reviewed in references^5-7^). Instruments automating volume-limited liquid handling extended nanoLC tandem HRMS to improving sensitivity. Up to 4,002 proteins were identified from yeast using LC-HRMS.^8^ Recent nanoPOTS (N2) arrays analyzing 100 pg of tryptic protein digest identified ~1,300 proteins from single murine cells.^9^ The automated single cell proteomics (SCoPE2) workflow enhanced throughput to ~200 single cells in 24 h using a refined nanoLC system, while reporting on ~1,000 proteins per cell.^10^ These experiments deepened the detectable coverage of the single cell proteome, albeit at the expense of long experimental times (~2–5 h per analysis).

Capillary electrophoresis (CE) HRMS delivers speed and sensitivity for single-cell analyses. We and others developed robust methodologies and custom-built CE-ESI instruments capable of zeptomole–femtomole sensitivity (reviewed in references ^11-12^). For example, ~200 protein groups were identified from 5 ng of *Pyrococcus furiosys* digest^13^ and up to 1,209 proteins groups from single embryonic cells dissected from^14-15^ or analyzed *in situ/vivo*^16-17^ in chordate embryos of important biological models (*Xenopus laevis*, zebrafish). A self-built CE platform and a sheath-flow tapered-tip CE-ESI design equipped HRMS with ~260 zmol sensitivity (156,000 copies) and robust operation on a quadrupole time-of-flight (TOF) mass spectrometer, allowing us to identify ~217 protein groups from ~1 ng protein (approximating ca. two neurons).^18^ An electrokinetically pumped low-flow sheath-flow CE-ESI interface achieved 330 zmol sensitivity, identifying 1,249 protein groups from 300 ng protein digest from *Xenopus laevis* eggs,^19^ ~4,400 protein groups from 220 ng of K562-cell digest via 2 h of separation,^20^ and ~100 proteins from 16 pg of *E. coli* protein digest.^21^ Proteins in these experiments were identified in a short amount of time, usually within an hour.

We recently showed that compressed peptide separation during electrophoresis challenges proteomics sensitivity.^4, 18^ Peptides in these experiments separated over a ~20–30 min effective separation window in total analysis times that were ~2–10-times faster than those typical in nanoLC. The ion flux in CE was chemically complex yet transient, lasting a few seconds only. These conditions challenged the duty cycle of peptide isolation, activation, and fragmentation due to limits in the duty cycle of modern tandem HRMS. In our recent work analyzing protein amounts estimating to ~2 mammalian neurons, only ~43% of peptide-like ion signals with unique m/z and separation time (called molecular features, MFs) were identifiable as peptide spectral matches (PSMs) from ~217 proteins.^4^ Supplementing electrophoresis with an orthogonal mode of separation, such as high-pH fractionation, allowed us to reduce the complexity of the chemical space in the temporal dimension, improving HRMS sensitivity to ~225 proteins from amounts approximating to single neurons.^4^ Even with orthogonal separation, however, the resulting MS/MS transitions exhausted the bandwidth of data-dependent acquisition (DDA) targeting the most abundant peptides for fragmentation. In mass spectrometers enhancing resolution in trade of speed, such as the orbitrap vs. TOF, we anticipate compact separation to exacerbate identifications in CE.

Iterative DDA presents a potential a way around the challenge. In nanoLC HRMS, the same sample is typically analyzed multiple times using the same DDA method (technical replicates) to leverage stochastic selection of complementary sets of peptides for identification.^22-24^ Triplicates detected ~20% more proteins, with this gain quickly diminishing after the 5^th^ replicate, reaching ~95% saturation by the 10^th^.^22^ Other DDA methods employed advanced iterative strategies to boost identifications by exclusion or inclusion of ions of interest (mass-to-charge, m/z values).^25-28^ Precursor ion exclusion (PIE), for example, progressively excluded peptides by dynamically updating the ion exclusion list for each replicate analysis. The approach helped nanoLC-HRMS to 533 proteins from 5 replicates of 1 μg of yeast whole cell lysate digest, equating to an ~51% improvement compared to the standard approach.^25^ Modified versions of PIE enhanced peptide identification by 52% with 4 measurements on 16 μg of IgG1 monoclonal antibody digest.^29^ Alternatively, post analysis data acquisition (PAnDA) iteratively prioritized unselected peptide features for targeted analyses with each replicate.^30^ The inclusion ion list measured 1,059 protein groups from 6 replicates of 4 μg of protein digest (*C. elegans*), resulting in an ~18% increase in identifications compared to standard DDA. By anticipating peptides in real-time based on a known elution order, 80% more proteins were identifiable from 4 measurements in mouse.^31^ These DDA approaches from studies employing nanoLC raise promising potentials to deepen proteomic sensitivity using CE-HRMS.

Here, we build on iterative DDA to advance ultrasensitive CE-HRMS to neuroproteomics on an orbitrap tandem mass spectrometer. DDA methods executing a nested series of ion exclusion lists formed rungs of our “DDA ladder,” allowing us to probe deeper into the proteome of cultured primary neurons. The method combined high spectral resolution from an orbitrap mass spectrometer (~120,000) with simplicity. Unlike earlier variants of DDA, we established a static list of ions for exclusion based on abundance tiers. In a proof of principle, the method achieved detection of 428 proteins (415 quantified), including many important in neurobiology, from sample amounts estimating to ~10 neurons. The DDA ladder expands the family of tandem MS approaches and emerging technologies, such as μCE-ESI, supporting ultrasensitive neuroscience research.

## METHODS

### Materials

Unless otherwise stated, all materials were purchased at reagent grade from Sigma-Aldrich (St. Louis, MO). Standard lysozyme, myoglobin, and papain dissociation system were obtained from Worthington Biochemical Corp (Lakewood, NJ). Sodium dodecyl sulfate (SDS, 10% v/v) was from Amresco (Solon, OH). Ethylenediamine tetraacetic acid (EDTA), LC-MS grade acetonitrile (ACN), formic acid (FA), acetic acid (AcOH), methanol (MeOH), MS-grade trypsin protease, and water (Optima) were supplied by Fisher Scientific (Fair Lawn, NJ). Ammonium bicarbonate was from Avantor (Center Valley, PA). Hank’s balanced salt solution, fetal bovine serum, penicillin-streptomycin, minimum essential medium, neurobasal medium, glutamine, and poly-L-ornithine were obtained from Gibco (Grand Island, NY). B27, N-2, glutamax, and pyruvate supplements were from Thermo Fisher Scientific (Waltham, MA). Fused silica capillaries (20/90 μm inner/outer diameter and 75/350 μm inner/outer diameter) were from Polymicro Technologies (Phoenix, AZ) and used as received. The stainless steel tapered-tip metal emitter (100/320 μm inner/outer diameter) was manufactured by New Objective (Woburn, MA). To minimize peptide loses due to nonspecific adsorption, the shot-gun proteomics workflow was conducted in LoBind protein microtubes from Eppendorf (Hauppage, NY).

### Buffers and Standard Solutions

The *cell lysis buffer* was prepared to contain: 5 mM EDTA, 20 mM Tris-HCl, 1% (v/v) SDS, and 35 mM NaCl. The *neuronal plating medium* was prepared as described elsewhere^32^ to contain: 0.6% (v/v) D-glucose, 10% (v/v) horse serum, 1% glutamine, and 1% penicillin-streptomycin in 1× MEM. The *neuro-maintain medium* was prepared to contain: 1% penicillin-streptomycin, 1% glutamax, 1% N-2, 2% B27, and 1% pyruvate in neurobasal medium. Prior to usage, all media were filtered through a 0.2 μm porous mesh.

### Neuron Culture

All procedures regarding the maintenance and humane treatment of mice were authorized by the Institutional Animal Care and Use Committee of the George Washington University (Approval Number A274). Adult pregnant C57BL/6 dams of mice (*Mus musculus*) were purchased from Charles River Laboratories (Wilmington, MA). Primary cultures of isolated mouse hippocampal neurons were prepared as described elsewhere.^32^ After 14 days of culture, each plated well was washed 3 times with 1 mL of SDS (10% v/v), and the cultured neurons were gently scraped to be transferred to a 1 mL LoBind microtube using micropipette with a LoBind tip. The collected neurons were centrifuged at 800 × *g* for 5 min at 4 °C and stored at −80 °C until processing for measurement.

### Bottom-up Proteomics

The collected cultured neurons were processed using a standard bottom-up proteomic workflow (see reference ^24^). 200 μL of lysis buffer was added to the collected neurons with 15 min of sonication in cold water bath (~4 °C). The lysate was reduced (5 μL of 1 M dithiothreitol, DTT, 30 min at 60 °C), alkylated (10 μL of 1 M iodoacetamide, IAD, 15 min in the dark) and quenched (5 μL of DTT). The lysate was centrifuged at 11,200 × *g* for 10 min at 4 °C, and the supernatant was transferred into a new 2 mL LoBind microtube. Proteins in the aliquot were purified by precipitation in 1 mL of chilled acetone (−20 °C, 5× volume of sample aliquot) over 12 h, followed by centrifugation at 11,200 × *g* for 10 min at 4 °C. The protein pellet was rinsed with chilled acetone, vacuum-dried at room temperature, and reconstituted in 200 μL of ammonium bicarbonate (50 mM). The final concentration of protein content was 0.5 μg/μL based on the standard bicinchoninic acid assay (Thermo Scientific, Waltham, MA). Aliquots of 40 μL were transferred into 5 separate 500 μL LoBind microtubes to serve as technical replicates. Each aliquot was digested with 0.8 μL of trypsin (1 mg/mL) over 12 h at 37 °C. The resulting peptides were vacuum-dried and reconstituted in 40 μL of 50% (v/v) ACN containing 0.05% (v/v) AcOH, chosen to help field-amplified sample stacking during electrophoresis.^18^ The peptide concentration of each aliquot was quantified using the Pierce colorimetric peptide assay (Thermo).

### Microanalytical CE-ESI-HRMS

This study utilized a home-built co-axial sheath-flow μCE-nanoESI platform^18^ with the following settings: CE, 90 cm capillary at 23 kV vs. Earth ground (applied to the inlet); CE-ESI sheath solution, 50% MeOH (0.1% FA) supplied at 300 nL/min; ESI, +2,700 V at 2 mm from the MS inlet, operated in the cone-jet spraying regime for efficient ion generation.^33^ Peptide ions were detected using a quadrupole orbitrap mass spectrometer (Q Exactive Plus, Thermo Scientific, Waltham, MA) between *m/z* 350–1,800 at 35,000 FWHM resolution (128 ms transient, 110 ms fill time). Higher-energy collisional dissociation (HCD) was triggered by DDA with the following instrumental settings: peak width (FWHM), 13 s; mass exclusion mass tolerance, 10 ppm; isolation window, 0.8 m/z; peptide matches, on; apex trigger, turned off; ion signals excluded below +2 charge state; ion signal intensity threshold, 1.5 ×10^3^ counts; maximum ion trap time, 50 ms for MS^1^. HCD employed the following conditions: m/z range, 200–2,000; resolution, 17,500 FWHM (64 ms transient, 50 ms fill time) for MS^2^; normalized collision energy, 32; dynamic exclusion, 5.0 s; maximum IT, 50 ms for MS^2^; TopN, 15; loop count, 15.

### Data Analysis

Using *MaxQuant*^34^ (version 1.6.17.0), the MS–MS/MS data were searched against the mouse proteome database containing 16,915 entries (downloaded from SwissProt on September 13^th^, 2017). The search parameters were: digestion, tryptic; missed cleavages, maximum 2; minimum peptide length, 5; minimum number of unique peptides, 1; fixed modification, carbamidomethylation at cysteine; variable modification, oxidation at methionine; match tolerance for main search peptide tolerance and MS/MS, ± 4.5 ppm and ± 20 ppm, respectively; isotope match tolerance, 2 ppm; decoy mode, revert; label-free quantitation, enabled; match between runs, turned off. Proteins were identified and filtered to <1% false discovery rate (FDR), computed against a reversed-sequence decoy database. The resulting proteins were grouped based on the closest parsimony principle. *Common contaminant proteins* were manually removed from the list of proteins that are reported in this work. MFs were surveyed in MzMine 2.0^35^ with the following parameters; mass detector, exact mass; noise level, 5,000; chromatogram builder, on; scans, MS^1^; min time span (seconds), 12 s; minimum height, 5,000; m/z tolerance, 0.05 m/z or 20 ppm. Gene ontology annotation of biological and molecular function were conducted in the PANTHER Classification System.^36^ Brain anatomy was modeled using Brain Explorer 2.^37^

### Safety Considerations

Metal tapered-tip emitters and fused silica capillaries pose needle-stick hazard and should be handled with care. Handling of chemicals followed standard safety protocols. Electrically conductive components of the μCE-ESI-HRMS platform were grounded (Earth) or isolated to eliminate electrical shock hazard.

## RESULTS AND DISCUSSION

### Addressing A Technological Gap

The goal of this study was to enhance CE-ESI-HRMS sensitivity for ultrasensitive neuroproteomics. CE-HRMS enabled the analysis of hundreds to thousands of different molecules in single cells and limited protein digests in exceptional sensitivity, in cases ~200–1,000-times higher than nanoLC.^4, 14-18^ Biomolecules in these experiments migrated in a considerably short time frame, typically with an effective separation window lasting ~15–30 min. For complex ‘omes, particularly an expanded peptidome in bottom-up proteomics, compressed separation taxes identification due to limitations in HRMS-MS/MS sensitivity and speed.^4^ To enhance detection, electrophoresis employing long separation times^38^ or orthogonal dimensions, such as ion exchange^39-40^, size-exclusion^41^, or high-pH C18 chromatography^18^ reduce molecular complexity over time by tailoring the transient ion flux to the duty cycle of MS/MS. Technological implementation, however, becomes increasingly challenging for trace amounts. Peak broadening due to longer separations is anticipated to reduce sensitivity, as modeled by the Van Deemter equation. Analyte transfer between separation dimensions risks peptide losses. These technologies also require skilled expertise and access to advanced, often custom-built, instrumentation, further constraining ultrasensitive CE-HRMS to specialized laboratories.

To advance CE sensitivity, we proposed to improve the success of data acquisition in HRMS–MS/MS. Our experimental rationale is shown in **Figure 1**. We used a custom-built (onedimensional) μCE-ESI system^18^ to analyze ~1 nL of protein digest from 250 nL of sample in each measurement.^4, 18^ These samples contained protein amounts approximating to ~2 neurons. Other custom-made or commercial CE systems with compatibility to handling limited sample volumes serve as alternative platforms in other laboratories. We reasoned that a microanalysis by CE allows for deepening proteomic coverage via repeated measurements if each replicate reports progressively deeper into the proteome, all the while still consuming trace amounts of total proteins. Our goal in this work was to analyze total protein amounts that are commensurate with limited populations of cells, such as less than 10 neurons.

**Figure 1.**
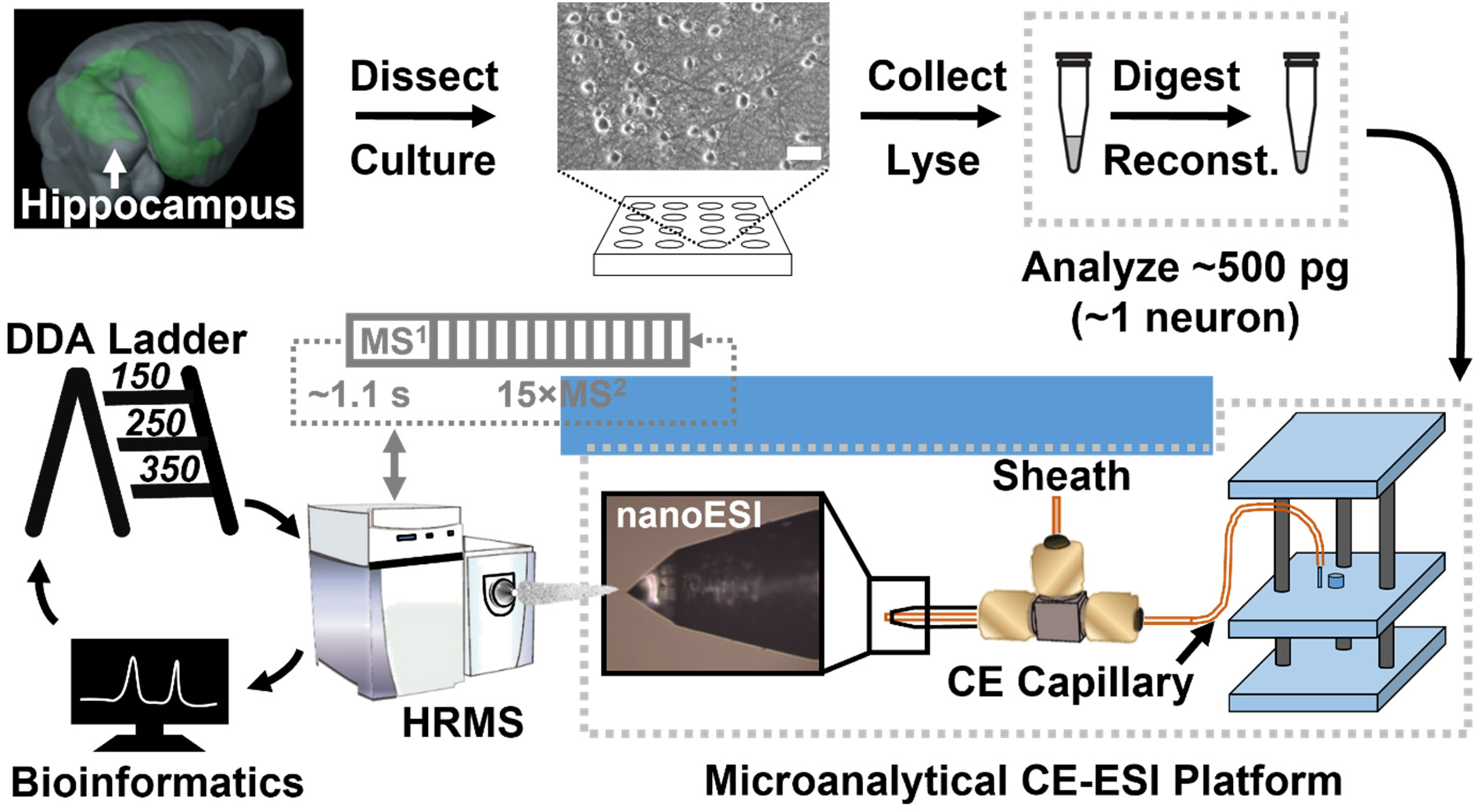
Experimental strategy of the nested DDA ladder for ultrasensitive CE-ESI-HRMS. The 150, 250, and 350 most abundant (top) peptides were excluded during replicate measurement of protein digests, with each replicate approximating to a single neuron (~500 pg/cell). Scale bar, 50 μm.

Method development began with control experiments optimizing the mass spectrometer for sensitivity. A protein digest was prepared from a culture of primary hippocampal neurons (mouse). The digest was diluted so ~1 ng of material was analyzed per replicate for method development, estimating to ~2 neurons.^4^ The peptides were electrophoresed, ionized, and detected in a quadrupole orbitrap tandem mass spectrometer (Q Exactive Plus, Thermo, see **Methods**). We chose DDA to govern MS/MS, because this method is robust, well-established, and available on most mass spectrometers. We anticipate these considerations to facilitate adoption of the method in other laboratories. A single-stage MS scan surveyed signals (m/z values), followed by fragmentation of the most abundant ions with a peptide-like isotope pattern. DDA parameters were optimized for identification based on technical duplicates. A dynamic exclusion of 5 s yielded 380 proteins (vs. 334 proteins at 2 s, 286 at 10 s, and 330 at 15 s), an MS^1^ automatic gain control (AGC) of 10^6^ counts returned 432 proteins (vs. 364 proteins at 5×10^5^ counts), and selection of the top 15 most abundant ions for MS/MS (TopN) gave 518 proteins (vs. 359 proteins at Top12, 474 at Top20). A list of proteins identified from the triplicates consuming a total of 3 ng of digest is tabulated in **Supplementary Table 1A** (**Table S1A**).

These measurements were tested for scalability to lesser protein amounts. Analysis of 500 pg of digest reported 190 proteins (~1 cell). The number of identifications increased with each technical replicate. Technical duplicates gave 213 proteins (~2 cells) and triplicates returned 244 proteins cumulatively (~3 neurons, see proteins in **Table S1B**). Detection of proteins with important roles in molecular pathways agreed with our earlier studies, demonstrating robust performance from CE-ESI-HRMS.^4, 18^ **Table S1C** uses the PANTHER Classification System to compare these pathway functions between 1 ng and 500 pg of measurements. Reduction of sample amounts resulted in missed detection for 339 proteins, or ~58% of all proteins identified. These proteins were implicated in several pathways important for health, including signaling. Neurobiological research would benefit from enhanced CE-HRMS sensitivity capable of deeper coverage of molecular pathways in tissue biopsies and small populations of cells.

To tackle this challenge, we began with understanding the current sources of limitations hindering DDA. In shot-gun proteomics, protein identification requires fragmentation of MFs that can be assigned to a peptide sequence with low error (<1% FDR), producing peptide spectral matches (PSMs). In our optimized method (see earlier), each DDA cycle targeted maximum 15 MFs. **Figure 2** compares the temporal evolution of MFs and PSMs to reflect on the utilization efficiency of the MS/MS duty cycle. During the ~60 min experiment, 935 peptides were detected (<1% FDR) migrating through the capillary within ~15 min, between ~30 min to ~45 min (**Fig. 2A**). This short effective separation window witnessed a rapidly increasing number of MFs. Their success of assignment as a PSM substantially varied. At the most compact zone of separation, only ~20–30% of MFs were identified. The success of identification increased to ~35–50% as molecular complexity relaxed toward the end of electrophoresis. These results from a quadrupole orbitrap mass spectrometer agreed with limited identification success (~45%) that we reported from a quadrupole time-of-flight instrument,^4^ highlighting an analyzer-independent need for better data acquisition during CE-HRMS. **Figure 2B** evaluates MS/MS transitions underpinning the PSMs on the orbitrap. In response to increasing MFs, DDA boosted the rate of MS/MS transitions, albeit only up to the minimal duty cycle. Notably, ~95.5% of the MS/MS transitions exhausted the maximal C-trap fill time. Therefore, DDA curtailed MS/MS rate to trade identification for sensitivity.

**Figure 2.**
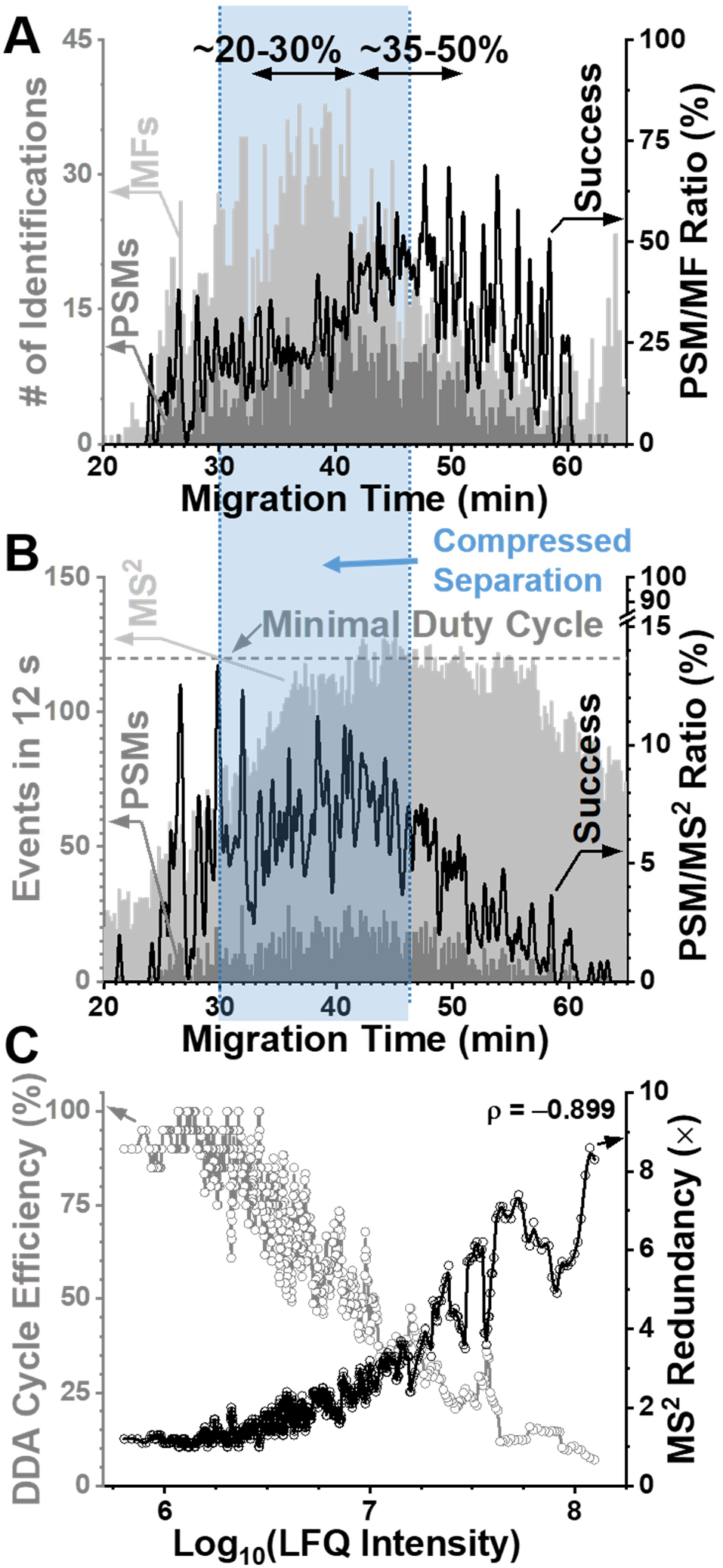
Challenges in trace-sensitive neuroproteomics using CE-HRMS with standard DDA. **(A)** Comparison of molecular features (MFs) and peptide spectral matches (PSMs), uncovering rapidly diminishing success at identifying putative peptide signals during the most compressed portion of electrophoresis. **(B)** Evaluation of tandem MS events underlying peptide identifications, showing optimal success at minimal MS/MS duty cycle. **(C)** More abundant peptides were fragmented at increasing redundancy (black curve), causing a steep decline in the efficiency of the DDA cycles identifying PSMs as peptides (10-point rolling average shown, white curve). Key: ρ, Coefficient of Pearson product-moment correlation.

But conducting many MS/MS scans does not necessarily yield complementary PSMs. In a representative dataset, 2,196 different PSMs were acquired from 1,203 different peptides; thus, the same peptide was fragmented ~1.8-times on average (MS^2^ redundancy). For example, the peptide HFFTVTDPR (from the protein Sideroflexin-3) was fragmented once, whereas the ~100-times more concentrated NLDIERPTYTNLNR (Tubulin alpha-1B chain) was fragmented 8 times. **Figure 2C** systematically compares MS^2^ redundancy with DDA efficiency as a function of peptide abundance using label-free quantification (LFQ).^15^ More abundant peptides were more likely to yield PSMs. This redundancy ca. tripled with each decade of ion signal abundance across the ~3 log-order observed dynamic range. In parallel, the efficiency of cycle utilization during DDA, approximated here as the PSMs/peptide ratio, took a steep decline. The datasets were strongly anticorrelated (ρ = −0.899, Pearson). Between the extremes, an ~9-fold increase in MS^2^ redundancy translated to a 15-fold reduction in cycle utilization (see **Fig. 2C**). Therefore, more abundant peptides were not only increasingly more likely to be redundantly fragmented, but they also required progressively more time from DDA, exacerbating a decline in identification success.

### A Guided Design for the DDA Ladder

These limitations, we reasoned, could be (partially) addressed by reducing PSM redundancy. To develop this method, we sought to better understand DDA economy. **Figure 3A** traces the cumulative for ion signal abundance and PSMs as a function of concentration (calculated LFQ intensities). Ca. 65%, ~75%, and ~85% of total signal abundance was produced by the 150, 250, and 350 most abundant (top) peptides. They respectively accounted for ~25%, ~37%, and ~50% of all the PSMs performed. This information supported our working hypothesis: DDA redundancy hindered protein identifications.

**Figure 3.**
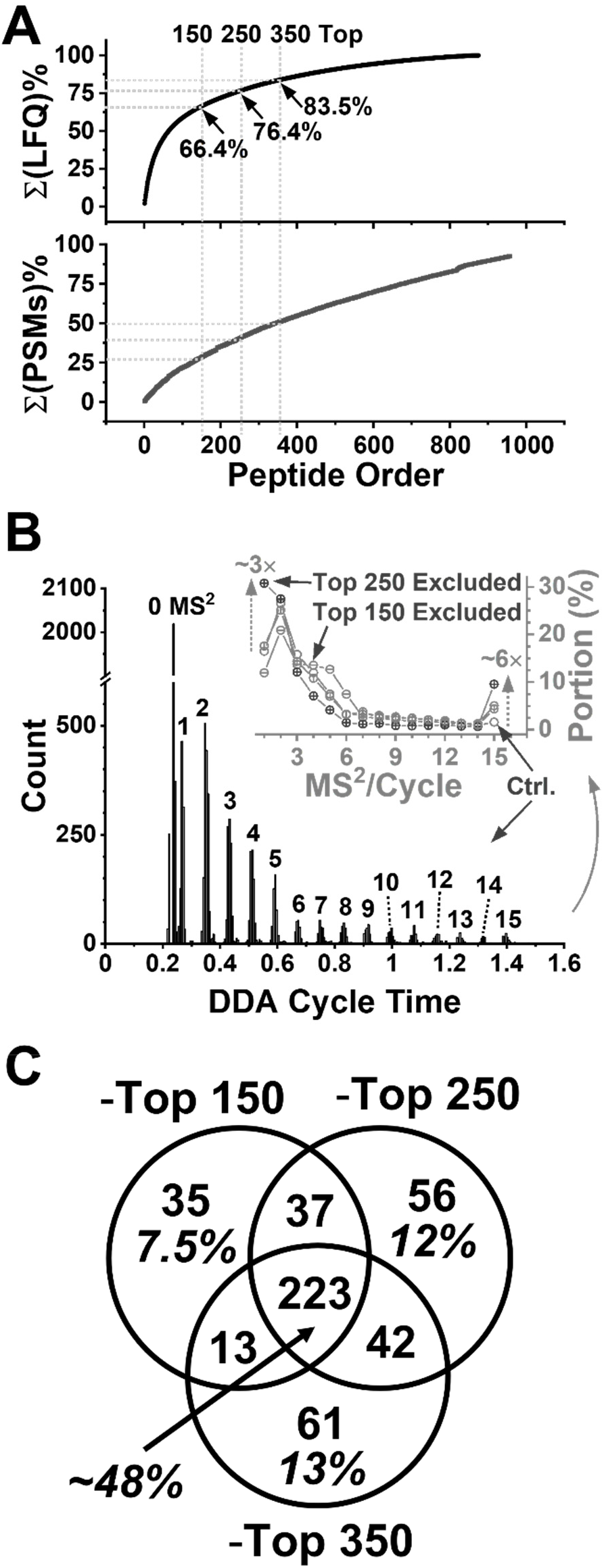
Assessment of MS–MS/MS economy to guide DDA ladder design. **(A)** The 150, 250, and 350 most abundant peptides accounted for the majority of signal abundance (LFQ intensity) and peptide spectral matches (PSMs) in control experiments. **(B)** Evaluation of the DDA cycle time, revealing underfilling of the top-15 peptide ions targeted for tandem MS. **(Inset)** Most DDA cycles were completed after fragmenting only 1–5 peptide ions. Exclusion up to the 250 most abundant ions (−250 Top) improved full utilization of the DDA cycle. **(C)** Each additional rung of the DDA ladder identified a complementary set of proteins in addition to the control (without ion exclusion).

After validating this hypothesis, we developed an improved version of DDA in this study. Based on our results, we proposed that exclusion of these redundant features during replicate analyses would free up DDA bandwidth to lower-abundance peptides in subsequent measurements. Rather than executing the conventional strategy, in which the most abundant peptide ions are excluded at the risk of missing corresponding protein identifications, our approach was to nest DDA methods based on signal abundance. In theory, this approach afforded detection for the abundant proteins based the preceding set of measurements (see triplicate later), before filtering them out in subsequent analyses. It followed that protein identifications from each abundance level of exclusion would mutually complement each other, with each exclusion level deepening quantitative depth. As proof of principle, we supplemented the control DDA (without ion exclusion) with three nested 3 DDA methods that progressively excluded ions based on ion signal abundance (recall **Fig. 1**). These DDA methods offered vantage points at different depths into the proteome, akin to rungs of a ladder; therefore, we termed the method a “DDA ladder.”

The DDA ladder was tested for performance. Protein amounts of 500 pg, which approximates to a single neuron (cell),^4^ were measured in triplicate as the control (without exclusion of ions). The resulting proteins were ranked by LFQ abundance (recall **Table S1B**). The triplicates were repeated while excluding the most abundant 150, 250, and 350 (“top”) ions, identifying 308 (**Table S1D**), 358 (**Table S1E**), and 339 (**Table S1F**) proteins, respectively. We inspected utilization of the DDA cycles by calculating the time that was spent between the survey scan and the last tandem MS event. **Figure 3B** exemplifies the analysis for conventional DDA and expends the analysis the experimental conditions (see **inset**). Ca. 42% of the cycles were completed without fragmentation (see 0 MS^2^); therefore, these scans did not identify any protein. Of the cycles performing tandem MS, ~10–30% targeted 1–5 peptide ions (see MS^2^/Cycle in the **inset**). Only less than ~2% of the cycles were filled with 15 MS^2^ scans. Disregarding the abundant ions provided improvement. By excluding the top 250 ions, the fraction of DDA cycles completing 1 tandem MS was tripled and those conducting the maximal 15 MS/MS scans was improved by 6-fold. Cycles completing transitions between these boundaries became less likely (see 2–-14 MS^2^ scans). Therefore, exclusion of the most abundant ions enhanced utilization of the limited MS/MS duty cycle.

Better MS/MS utilization raised the possibility of identifying complementary proteins. **Figure 3C** compares identified proteins from excluding the top 150, 250, and 350 peptide ions. While ~48% of proteins were commonly identifiable, each of the tested rungs reported on a different set of proteins. Each tested exclusion level provided ~4.5–13% proteins that were only identifiable at each specific DDA rung. **Table S1G** shows pathway enrichment analysis for these proteins (PANTHER). Cytoskeletal regulation, Huntington disease, gonadotropin-releasing hormone receptor pathways, Parkinson disease, glycolysis, and EGF and FGF signaling, Wnt signaling, and the TCA cycle were top pathways represented by proteins that were commonly identified at each rung of the ladder. These pathways were complemented with additional proteins from exclusion of the top 150 ions. The second rung of exclusion expanded pathway coverage with angiogenesis, endothelin signaling, and PDGF signaling. Exclusion of the top 350 ion signals provided additional proteins in the pathways covered by the lower rungs and may be beneficial if additional protein identifications are desired in a particular study.

The DDA ladder was benchmarked (**Fig. 4**). As the reference, standard DDA (without ion exclusion) was chosen, which is the closest neighboring technology to the DDA ladder presented here. **Figure 4A** predicts the cumulative number of proteins on the basis of up to quintuplet technical replicates (Control). As also found by others,^22^ rapid saturation in identification provided diminishing returns from the replicates. While triplicates identified 244 proteins in our study, the projections were 287 proteins at 6, 319 at 9, and 338 at 12 replicates. Each additional replicate after triplicate measurement returned only ~5% more proteins. Triplicates were a satisfactory trade between identification number, sample consumption, and analysis for our study. Each additional rung of the DDA ladder identified ~21%, ~34%, and ~42% more proteins compared to the control, improving cumulative identifications by ~42%, 24%, and 12%, respectively. With a diminishing return in cumulative identification, we chose the first two rungs (top 150 and 250 ions excluded) sufficient to supplement the control experiments for this study. Consuming ~500 pg proteins per analysis, the finalized DDA ladder for this study therefore required the analysis of a total of ~4.5 ng of protein between 9 measurements, estimating to proteins from ~10 neurons.

**Figure 4.**
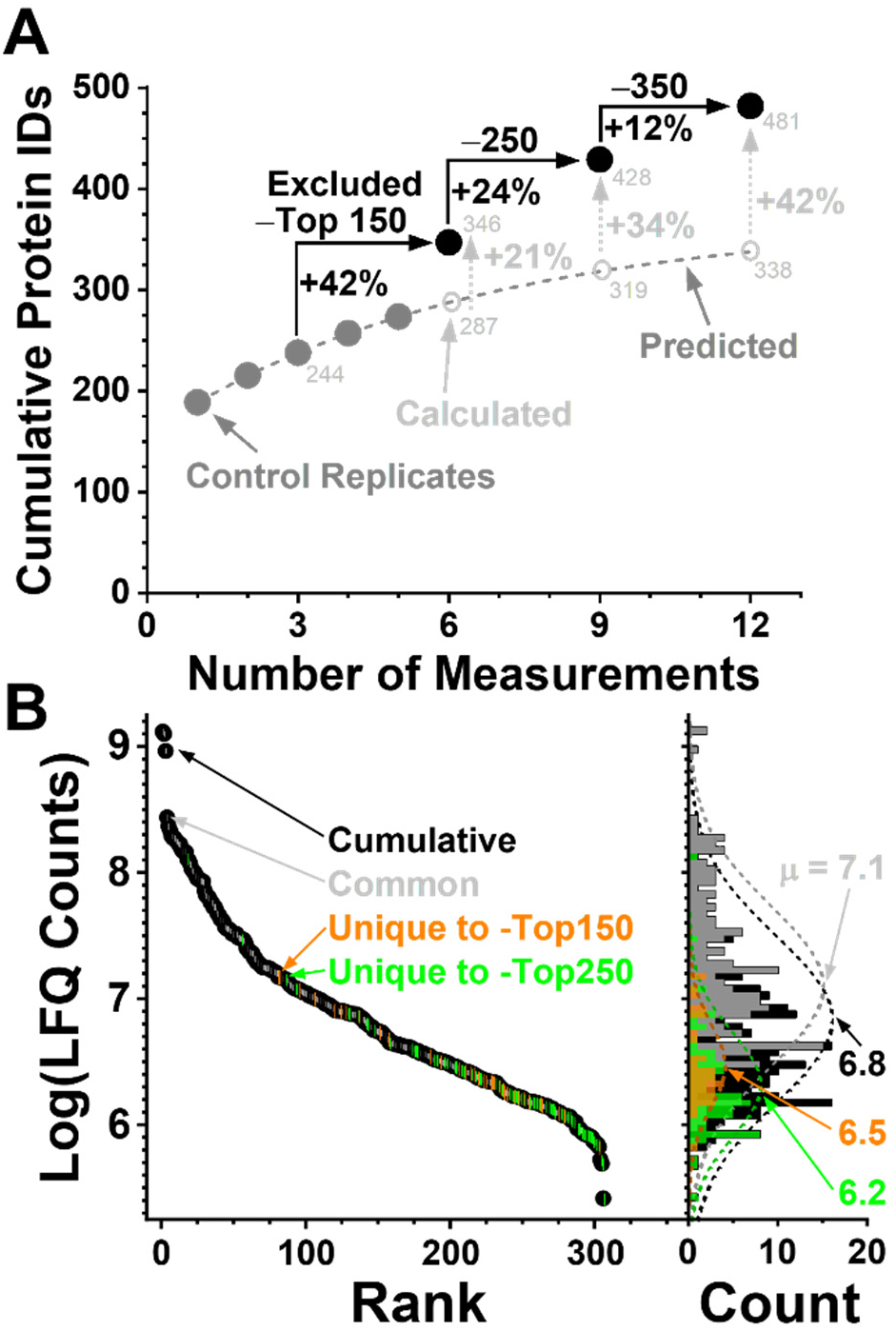
Depth of proteome coverage. **(B)** Cumulative protein identifications from technical replicates by standard DDA (Control) and a nested DDA ladder excluding the top 150, 250, and 350 ions. The first two rungs of the ladder were chosen sufficient for this study. **(B)** Dynamic range of quantification by the DDA ladder, revealing quantification of lower-abundance proteins with each tested rung. Proteins only quantifiable in the control (grey) and after exclusion of the 150 (orange) and 250 (green) most abundant peptides are overlayed on the cumulative (black).

We inquired about the resulting depth of proteome coverage. The DDA ladder identified 428 proteins, 415 of which were also quantified (see **Table S1G**). To compare the abundance of the quantified proteins, we minimized technical variability in LFQ between replicates. The LFQ values were scaled using linear regression on the basis of the commonly quantified proteins (control vs. top 150 excluded: R^2^ = 0.93; control vs. top 250 excluded: R^2^ = 0.90). **Figure 4B** evaluates the quantitative dynamic range of the proteins. The most abundant proteins were quantified without ion exclusion (see control with μ = 7.1, Gaussian mean). Exclusion of the top 150 and 250 ions expanded the linear dynamic range of quantification. Therefore, each additional rung of the DDA ladder helped us quantify proteins deeper in the concentration range occupied by the neuroproteome, supporting out working hypothesis.

Last, we asked whether deeper quantification also translated into better coverage of molecular pathways. We compared pathways that were enriched by proteins detected by standard DDA consuming ~3 ng (see proteins in **Table S1A**) and ~1.5 ng (**Table S1B**) as well as the DDA ladder using ~4.5 ng (**Table S1H**) of protein digest. The analysis results are shown in **Figure 5**. A list of the respective pathways is tabulated in **Tables S1I (Eye)**. The DDA ladder reported on all the pathways that were represented by analysis of 1.5 ng of digest using conventional DDA. Our method also recovered proteins from ~9 of 21 different pathways that were only detected by 3 ng of analysis previously. These pathways included angiogenesis, Alzheimer disease-amyloid secretase pathway, and several signaling pathways including alpha adrenergic receptor, toll receptor, PDGF, interleukin, and endothelin signaling. Further, the DDA ladder also uncovered previously unrepresented pathways, including transcription regulation by bZIP transcription factor, vitamin D metabolism and pathway, Hedgehog signaling pathway, sulfate assimilation, and cholesterol biosynthesis. Overall, our findings show that protein detection in a single cell-equivalent volume can be extended to identify proteins beyond the most abundant cytoskeletal and synaptic components to further probe signaling and metabolism in healthy and diseased cells.

**Figure 5.**
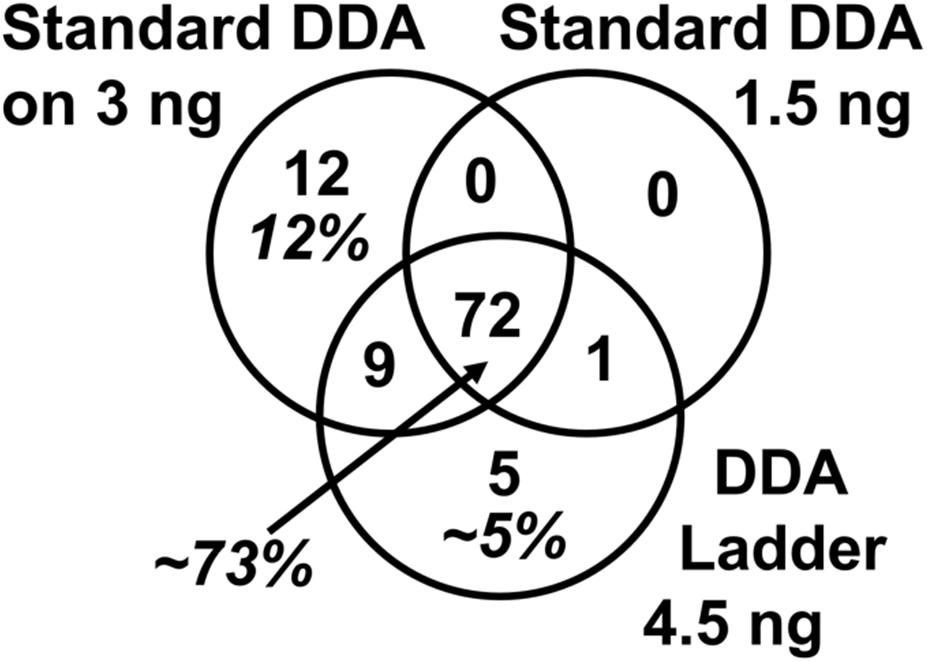
Gene anthology annotation of pathway coverage between the conventional DDA (control) and the DDA ladder. The numbers count different pathways enriched by the detected proteins. The DDA ladder uncovered several important pathways for neurobiology that would have been lost to limited utilization efficiency of the DDA cycle. Pathway annotation is provided in Table S1I.

## CONCLUSIONS

This study advanced CE-HRMS to trace-sensitive neuroproteomics. A DDA ladder encompassing two tiers of filters reduced a measurement bias toward high-abundance peptide ions, thus expanding the quantifiable linear dynamic range of the proteome. The method improved utilization of the limited DDA duty cycle, reduced redundant MS/MS events, and promoted the selection of a larger number of ions for targeted for fragmentation. Compared to standard DDA, the approach was better prepared to handle high peptidome complexity transiently unfolding during compressed electrophoresis. CE-HRMS employing the DDA ladder complements classical nanoLC-based proteomics with fast analysis (~30–45-min separation), ultrahigh sensitivity, and robust operation. Recent commercialization of CE instruments capable of handling limited amounts of samples, such as less than ~1–5 μL, may substitute the custom-built microanalytical CE platform in this study. With well-established usage and availability on commercial mass spectrometers, we anticipate the DDA ladder to be readily adaptable in other laboratories to other types or limited amounts of proteins via shot-gun proteomics.

Continuous advances at multiple stages of the proteomic workflow raise the possibility of further sensitivity improvements. Technologies enabling the isolation and handling of miniscule amounts of proteins with reduced loss, for example, by *in vivo* microsampling^16^, nanoPOTS^9^, and ScoPE^10^ offer viable solutions for biopsies and limited populations of cells, including single cells. Detection of 428 proteins from protein amounts estimating ~10 neurons in this study, including many with important biological functions in homeostasis and disease, marks another technological leap in ultrasensitive proteomics, expanding the bioanalytical toolbox of neuroscience.

## ACKNOWLEDGEMENT

This work was supported by the Arnold and Mabel Beckman Foundation Beckman Young Investigator Award (to P.N.) and a COSMOS Club Award (to S.B.C).

## Author Contributions

P.N. and S.B.C. designed the research. P.M-L. and M.C.M. prepared the neuron culture. S.B.C. processed the neuron culture, prepared the sample, and performed the measurements. S.B.C. and P.N. analyzed the data and interpreted the results. P.N. and S.B.C. wrote the manuscript: All authors commented on the manuscript.

